# RSVdb: A comprehensive database of transcriptome RNA structure

**DOI:** 10.1101/710236

**Authors:** Haopeng Yu, Yi Zhang, Qing Sun, Huijie Gao, Shiheng Tao

**Affiliations:** College of Life Sciences and State Key Laboratory of Crop Stress Biology in Arid Areas, Northwest A&F University, Yangling, Shaanxi, 712100, China; Bioinformatics Center, Northwest A&F University, Yangling, Shaanxi, 712100, China; College of Animal Science and Technology, Northwest A&F University, Yangling, Shaanxi, 712100, China

**Author notes:** These authors contributed equally to the paper as first authors (Haopeng Yu, Yi Zhang).

## Abstract

RNA fulfills a crucial regulatory role in cells by folding into a complex RNA structure. To date, a chemical compound, dimethyl sulfate (DMS), has been developed to effectively probe the RNA structure at the transcriptome level. We proposed a database, RSVdb (https://taolab.nwafu.edu.cn/rsvdb/), for the browsing and visualization of transcriptome RNA structures. RSVdb, including 626,225 RNAs with validated DMS reactivity from 178 samples in 8 species, supports four main functions: information retrieval, research overview, structure prediction, and resource download. Users can search for species, studies, transcripts and genes of interest; browse the quality control of sequencing data and statistical charts of RNA structure information; preview and perform online prediction of RNA structures *in silico* and under DMS restraint of different experimental treatments; and download RNA structure data for species and studies. Together, RSVdb provides a reference for RNA structure and will support future research on the function of RNA structure at the transcriptome level.

## INTRODUCTION

Intracellular RNA is not only the carrier of genetic information but also plays a critical regulatory role by folding into complex RNA structures. RNA structure *in vivo* is now known to be involved in RNA transcription, splicing, and subcellular localization and affects the initiation of translation, the ribosome translation rate, and protein co-translation folding [1–6]. Several compounds have been developed to probe RNA structure *in vivo*, among which dimethyl sulfate (DMS) reagents were first used to detect global RNA structure in the transcriptomes of *A. thaliana*, *H. sapiens* and *S. cerevisiae* [7,8]. The methyl group of DMS can be donated to the hydrogen bonds of adenine, cytosine, and guanine residues (m^1^A and m^3^C), and base-pairing will reduce the reactivity of this reaction [9]. The exact location of m^1^A and m^3^C in the transcriptome can be resolved by the truncation of reverse transcription method (RT-stop method) or increased mismatch ratio method (MaP method), which can be applied to determine RNA structures *in vitro* and *in vivo* [10,11]. After combining with high-throughput sequencing, these methods can effectively probe the structure of RNA at the transcriptome level, allowing for further study of the function of RNA structure in cells [12–18].

We collected ten studies probing the transcriptome RNA structure by DMS since the method was first proposed and presented a comprehensive, visual, and user-friendly database, RSVdb (https://taolab.nwsuaf.edu.cn/rsvdb/). RSVdb not only collects and displays transcriptome RNA structures but also subdivides relevant studies into experimental samples with detailed statistics and provides highly customized RNA structure prediction, enabling users to make sufficient use of transcriptome RNA structure data. RSVdb includes 626,225 RNAs with valid DMS reactivity from 178 samples of 10 studies in 8 species (Figure 1B, Table 1). The database provides interactive charts showing statistics at the transcriptome level, sample level, and transcript level, including quality control of sequencing, read mapping statistics, the percentage of each base, the RPKM, the Gini coefficient, and statistics of DMS reactivity. Further, we provide prediction and display of the RNA structure under different conditions. Users can search for transcripts of interest and predict and visualize mRNA structures with DMS reactivity restriction of complete or partial sequences. Additionally, we provide a detailed user manual and download of DMS reactivity data. All charts in the database are interactive and can be exported online as figures or in text format, and all search boxes in the database support search suggestions. In summary, RSVdb makes it easy to browse, search, and visualize transcriptome RNA structure data.

**Figure 1.**
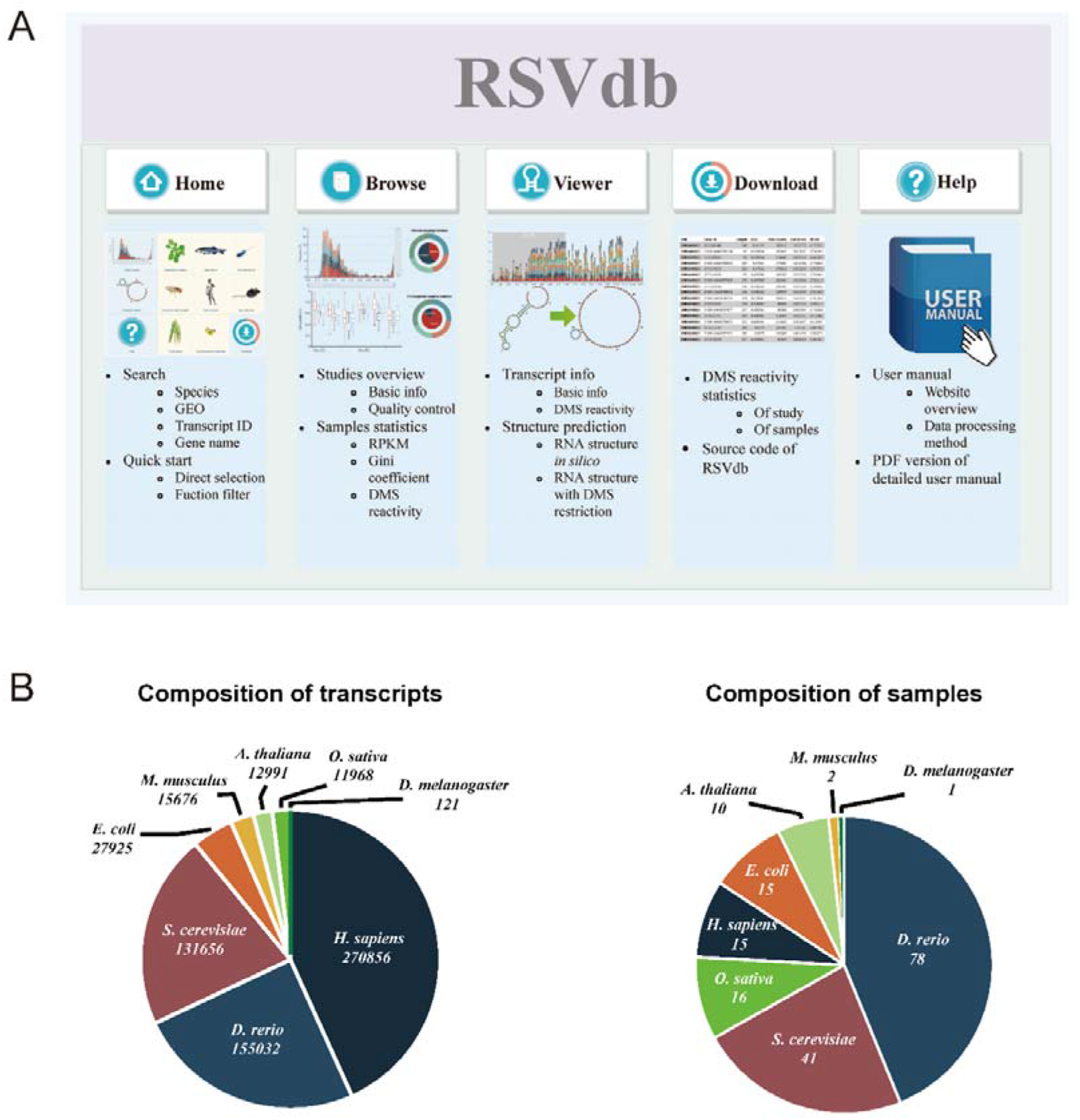
Schematic overview and main contents of RSVdb. (A) Website frame and central functions. (B) The composition of transcripts and samples among species.

## MATERIAL AND METHODS

### Data sources

The raw sequencing data were collected from Gene Expression Omnibus (GEO) and the Short Read Archive (SRA) databases. The current version contains 178 samples from 10 studies in 8 species: *A. thaliana, D. melanogaster, E. coli, H. sapiens, M. musculus, O. sativa, S. cerevisiae*, and *D. rerio.* DMS-seq, structure-seq, structure-seq2, and CIRS-seq methods use DMS modifications detected by reverse transcription (RT) halt sites, and these methods are collectively referred to as ‘RT-stop,’ including 151 samples from 5 studies of 6 species. DMS-MaPseq method uses reverse transcriptase mismatch instead of truncation products, referred to as ‘mutation profiling (MaP),’ including 27 samples from 4 studies of 5 species. The sequencing data of these samples were mapped to the reference genome and transcriptome, then normalized through data processing to screen out 622,429 RNAs with valid data.

### Sequencing data preprocessing

All SRA files were obtained from the SRA database of NCBI and converted to Fastq format by using pfastq-dump (https://github.com/inutano/pfastq-dump) and sratoolkit v2.9.6 (https://trace.ncbi.nlm.nih.gov/Traces/sra/sra.cgi?view=software). Quality control checks on raw sequence data were assessed by FastQC v0.11.8 (http://www.bioinformatics.babraham.ac.uk/projects/download.html). Multiple reports from a study generated by FastQC were combined into one report using MultiQC v1.8 (https://multiqc.info/). For Fastq files, adapters of each read were removed by Cutadapt v1.18 (https://cutadapt.readthedocs.io/en/stable/index.html) with parameters ‘-j 8 -n [N] -b’. All reads were filtered by Fastx_toolkit v0.0.14 (http://hannonlab.cshl.edu/fastx_toolkit/download.html) with a quality score higher than 30, and a read length greater than 21. Transcriptome sequences for eight species were downloaded from Ensembl, with each gene retaining only the longest transcript.

### Mapping to transcriptomes

Previous studies described methods to map reads to the genome and then extract transcriptome regions, and there were also methods to map directly to the transcriptome. We mapped raw reads to both the genome and the transcriptome and found that although there were more reads with the successful alignment of the genome, such parts that were more than the transcriptome tended to be located in the non-transcribed regions. Meanwhile, since the screening of reads requires the selection of unique reads, and the genome is prone to more multiple alignments, the number of valid reads mapped to the genome is reduced. Thus, by comparing genome mapping statistics, we recommend direct mapping to the transcriptome.

Methods for detecting transcriptome RNA structure using DMS can be roughly divided into two categories: the reverse transcription stop method (RT-stop method) and the increased mismatch ratio method (MaP method) using reverse transcriptase mutation. We mapped the data to the corresponding transcriptome. We used different strategies to map the two types of data to the transcriptome (Figure 2).

**Figure 2.**
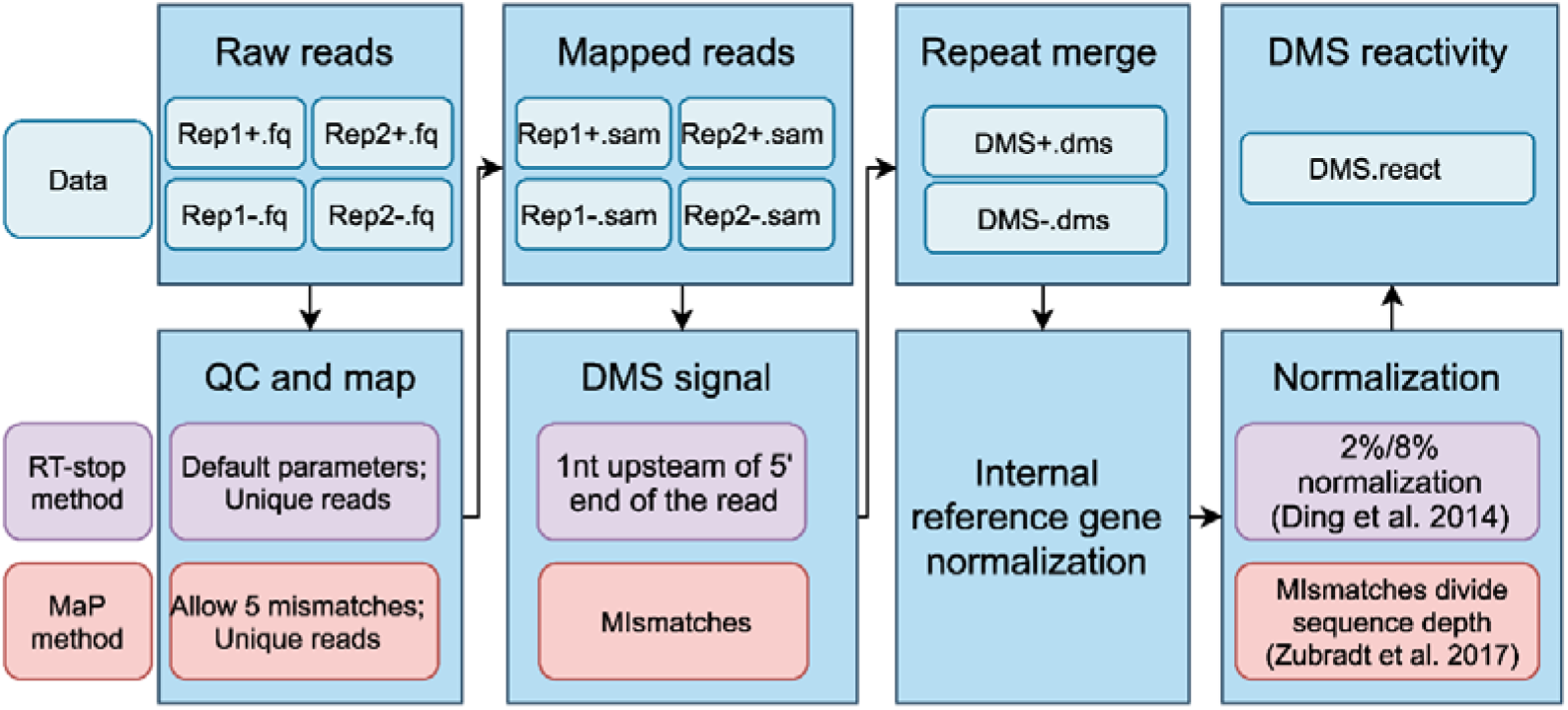
Transcriptome RNA structure data processing pipeline of RT-stop and MaP methods.

For the strategy involving RT-stop data processing, we mainly refer to the method of Ding et al. [19]. Hisat2 v2.1.0 (https://ccb.jhu.edu/software/hisat2/index.shtml) was used for sequence alignment with the following settings: hisat2 -p 16 -x reference_genome.file -U fastq.file -S SAM.file, where two mismatches were allowed. Only unique reads that did not have a second match were selected in the SAM file, while the reads mapped to the reverse strand or with first base mismatch were also removed. The quality control report of sequencing data from studies is displayed on the website. The actual truncated site is the nucleotide that is 1 nt upstream of the 5’ end of the mapped read. The repeated experiments were merged, and the internal parameter was selected by calculating the rRNA Pearson correlation coefficient of the combined samples and choosing the rRNA with the highest correlation (if all rRNA correlation coefficients were less than 0.95, other housekeeping genes were considered). On the “Browse” page of our website, we present the information on the internal reference gene and its correlation report for each study. Then, the samples were normalized with the DMS signal of the internal reference gene. Thus, the DMS reactivity between samples could be compared. We performed 2%/8% normalization on DMS signals [20]. In this step, the top 2% of DMS signal values are treated as outliers, and each DMS signal value (including the top 2%) divides the average of the remaining top 8% of DMS signal values to get normalized structural reactivity. Transcripts with no DMS signal in the 2% to 8% range were removed.

For the strategy involving MaP data processing, we mainly refer to the method of Zubradt et al. [21]. Tophat2 v2.1.1 with bowtie2 was used for sequence alignment with the following settings: – no-novel-juncs -N 5–read-gap-length 7–read-edit-dist 7–max-insertion-length 5–max-deletion-length 5 -g 3, where five mismatches were allowed. The mapping results were sorted by samtools v1.9 (http://www.htslib.org/download/), and all multi-mapping reads were removed from the BAM file. The mismatches of aligned reads to the reference sequence were summarized by using samtools mpileup with the following settings: samtools mpileup -f reference_genome.file -Q 15 -q 20 bam.file -o output.pileup. Then the mismatch ratio of the DMS signal was calculated for each adenine (A) and cytosine (C) nucleotide as mismatch numbers divided by the sequencing depth from the pileup file in a Perl script (removing indels). The strategy by which the biological replicates are merged, and the internal reference gene is normalized is the same as the method in the RT-stop section.

### DMS signal statistics

We used Perl scripts to conduct statistics for different stages of data processing, including the statistics after reads mapping after samples were merged and DMS signals were corrected by min-max normalized internal parameters and the statistics after DMS signal normalization.

After reads mapping, three types of statistics were requested, including (i) the proportion of mapped and unmapped reads of each study, (ii) the ratio of four bases, A, C, G, and U, to the DMS signal and (iii) the RPKM top 100 transcripts and RPKM distribution. RPKM was used as an index to measure the abundance of DMS signals for each transcript, which is defined below.

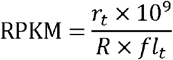

Where *r*_*t*_ is the raw DMS counts in a transcript, *R* is the total mapped reads in all transcripts, and *fl*_*t*_ is the feature-length of a transcript.

After samples were merged and internal parameters were normalized, the Gini coefficient of each transcript was determined. The Gini coefficient was used as an index to measure the strength of the RNA structure for each transcript, which was calculated by a sliding window, with a size of 50 A and C bases (G and T bases were removed) [8,22]. The Gini coefficient is defined as follows.

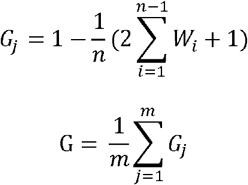

Where *w*_*i*_ is the cumulative DMS counts from the first base to the ith base as a percentage of the total DMS counts in the window *j*. n is the total number of A’s and C’s in a window, which is 50 here. *G*_*j*_ is the Gini coefficient of the *j*th window, and *m* is the total window counts of the transcript.

After the DMS signal was normalized, the data could be used to evaluate RNA structure, named DMS reactivity. We calculated the distribution of DMS reactivity for each sample and the top 100 transcripts. These statistics are presented in interactive charts on the website and are available for download.

### RNA structure prediction

The “Fold” in the “RNAstructure” package is adopted for RNA structure prediction [23,24] with the parameter ‘-dms.’ In the RNA structure prediction settings, we also added two customized options, “Threshold” and “Adjust.” “Threshold” is set to calibrate the DMS reactivity higher than the threshold to prevent deviation caused by excessive DMS reaction [19]. If the DMS reactivity is higher than the threshold, the reactivity of the base is assigned to the threshold. We found that different normalized strategies influence the prediction of an RNA structure, so we added the option “Adjust.” “Adjust” can modify the whole DMS reactivity of the RNA region (DMS reactivities multiply “Adjust parameter”), mainly for the studies without the DMS control sample. These two options are not enabled by default, and researchers can manage them if necessary.

### Overlapping bases ratio

Three species *E. coli*, *H. sapiens*, and *S. cerevisiae* had data from both RT-stop and MaP methods. Screening strategies are proposed to obtain comparable transcripts: data coverage should greater than 50%, and the relative deviation of the coverage of the two methods should less than 10%. Different samples of the same species are merged and proposed further statistics.

### Front-end interface and back-end structure

The implementation of the RSVdb database can be divided into two parts: front end (client-side) and back end (server-side). The back-end part is implemented by the python-flask web framework, which is responsible for server-side website operation logic and necessary data processing, and SQLalchemy, which is responsible for data storage and query. The front-end interface is written in HTML5, CSS, and Javascript and uses AJAX to interact with data on the server-side asynchronously. The front-end and back-end code of RSVdb are freely available via the GitHub repository (https://github.com/atlasbioinfo/RSVdb/).

## RESULTS

### Comparison between RSVdb and several other RNA structure databases

We compared the functions and features of RSVdb with several other RNA structure databases, including data types, species covered, RNA structure data presentation, and availability of RNA structure predictions.

As an earlier database of an RNA structure, RMDB contains a series of experimental data for single-nucleotide-resolution RNA structures generated by high-throughput sequencing techniques, describing 1,190 experimental data points of 145,447 RNAs (till Feb. 2020) [25]. The database provides search, visualization, and analysis capabilities for all users, structure viewer, and prediction function for registered users. However, RMDB has limited support for transcriptome-level RNA structure data and mainly including SHAPE data for *in vitro* measurement of the specific RNA structure. With the development of experimental techniques over the past few years, transcriptome-wide data have increasingly emerged in the analysis of RNA structures. In view of this, Structure Surfer, a transcriptome-based database, was proposed. It included *in vivo* (DMS-seq) and *in vitro* (PARS, ds/ ssRNA-seq) RNA structure data of *H. sapiens*, as well as *in vivo* and *in vitro* icSHAPE data of *M. musculus* [26]. The database provides searching and RNA structure profile visualization of two species and two studies but does not provide RNA structure prediction. The FoldAtlas database contains only complete transcriptome-wide high-depth structure-seq data from *A. thaliana* and provides circle plots and topology diagram visualization, *in silico* and *in vivo* RNA structure predictions and comparison. It also provides normalized reactivities as text files for download [27].

RSVdb is designed for visualization and prediction of transcriptome RNA structure. It aims to enable users to make sufficient use of transcriptome RNA structure data. The current version of RSVdb contains almost all transcriptome RNA structure data probed by DMS reagents, both RT-stop and MaP-related methods, including DMS-seq, structure-seq, structure-seq2, CIRS-seq, and DMS-MaPseq. Firstly, general statistics and an overview of these studies are shown on the “Browse” page. We analyzed these data at the transcriptome level, sample level, and transcript level to obtain data quality statistics, mapping ratio, A/C ratio, RPKM, DMS reactivities, and the Gini coefficient of different experimental samples. More importantly, RSVdb provides RNA structure prediction with highly customizable parameters and can be compared in different experimental conditions on the “Viewer” page. In addition, RSVdb provides detailed user manuals and download services. The recommendation of the RSVdb database can supplement the limitations of existing databases and provides a straightforward approach for researchers to find valuable information on transcriptome RNA structures.

### Browse RNA structure studies

The “Browse” page contains six parts, including the selection of species and studies, the detailed information of research, sequencing mapping, samples merged and data normalization, and the statistics of the RPKM, Gini coefficient, and DMS reactivity. Users can browse through these sections in order or jump from the left navigation bar to the corresponding section. The “Selection” section contains the selection of species and studies. After clicking, the information in other parts of this page would be updated asynchronously. The “Study info” section includes basic information on the study, sample, experimental processing, and sequencing data. The “Mapping” section includes the HTML version of the sequencing data quality control report, the sequence mapping and base composition statistics of the SRR data, and the transcripts of the RPKM top 100. The “Data merge” section shows the process of sample merging (the biological duplication is merged) and the distribution of the RPKM of combined samples. In the “Normalization” section, data were normalized by the internal reference gene. This part includes reports on the selected reference gene and its correlation coefficients, as well as the box plots of the Gini coefficient for different thresholds. Of course, we use different strategies to process and standardize RT-stop and MaP data (refer to the Methods section). The data in “RNA structure info” is further normalized. For RT-stop data, 2%-8% normalization is adopted [20], and MaP data is used to perform another normalization method [15]. In order to facilitate retrieval and database storage, we simplified the sample names in the original studies and showed the corresponding simplified sample names on the web page. In this section, we refer to the normalized data as “DMS reactivity” and show the top 100 transcripts of DMS reactivity and the DMS reactivity distribution of different samples. On the “Browse” page, we detail the sequencing data processing for the RNA structure studies. All diagrams are updated asynchronously and interactively, and images are available for download.

### Predict RNA structure with DMS restriction

In addition to viewing statistics charts from RNA structure research studies, RSVdb also supports RNA structure prediction with DMS restriction. After data screening, RSVdb currently contains 626,225 RNAs with valid DMS reactivities for RNA structural prediction. Users can search the gene name or transcript number of interest at the home page or “Viewer” page. The search box supports the search suggestions function, which will recommend relevant genes based on user input.

The “Viewer” page also contains six sections: species and research selection, gene search, genetic information, DMS reactivity display, RNA prediction settings, and RNA structure prediction (Figure 1A). After selecting the species and study, the user can locate the gene of interest through the search box and search suggestions. Then, the basic information and DMS reactivity of the target transcript in different samples is displayed in the “Gene Info” section. Meanwhile, the DMS reactivity of each base of a transcript is shown in the bar plot in the “DMS reactivity viewer” section. Users can drag the slider or directly input to select the RNA region, and in the “RNA structure prediction settings” section, the selected sequence and DMS reactivity are displayed. Finally, in the section “RNA structure prediction,” the RNA structure of the direct prediction of the RNA subsequence and the prediction after DMS restriction are shown.

We proposed a low minimum free energy region in the APA1 gene of *S. cerevisiae* as an example to demonstrate RNA structures under different experimental conditions using RSVdb. Two low free energy regions, S1 and S2, in the APA1 gene, could be obtained by the sliding windows method (Figure 3A). We performed research selection, gene searching, DMS reactivity display, and RNA structure prediction with DMS restriction on the “Viewer” page. The result revealed that the RNA structure of the S1 region weakens with increased temperature *in vitro* and has a weak structure *in vivo*. We noted that although the internal reference gene has normalized DMS reactivity in RSVdb, we provide “Adjust” and “Threshold” parameters in the “Prediction settings” (disabled by default), which can be adjusted to change the exhibited RNA structural prediction.

**Figure 3.**
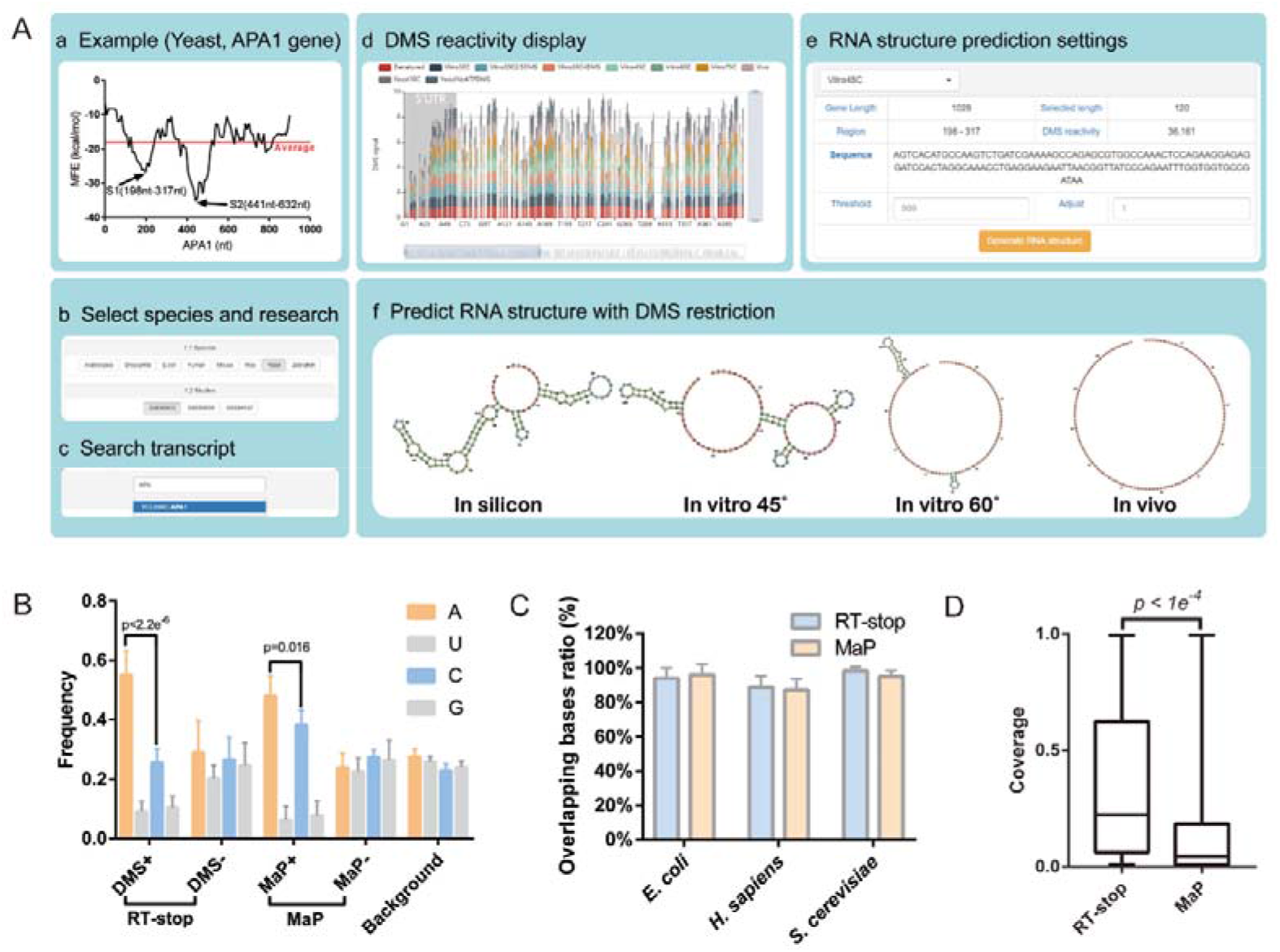
Predict and compare the RNA structure of target transcripts under several experimental treatments. (A) The processing pipeline of RNA structure prediction was presented as an example, the APA1 gene in Yeast. (B) The frequency of the four bases. (C) The ratio of overlapping bases of RT-stop and MaP methods. (D) Data coverage for the two methods.

### RT-Stop and MaP methods

DMS can methylate non-Watson-Crick base-paired adenines and cytosines (m1A and m3C), and this information can parse out single-stranded RNA sites and then resolve the conformation of mRNA structures *in vivo* while combining RNA structure models and algorithms [28–30]. Two main strategies are operated to detect the single-strand position of DMS to probe RNA structures may give rise to modification, namely, “RT-stop” strategy, which is suspended at one nucleotide upstream of the truncation base during reverse transcription, and “MaP” strategy, which is used to detect the mutation at the modification site [10,21]. However, the utilization of DMS to probe RNA structures may arise some limitations involving only A and C bases probed, other methylation mechanisms in the cell, the misincorporation of RT enzymes and modification block of RNA-binding proteins [31,32]. From the data perspective, we counted the A/U/C/G ratio both in RT-stop and MaP studies and found apparent bias of A sites [33], which revealed more obvious in RT-stop studies (Figure 3B).Although the G and U sites were removed in a further calculation, at least 5.57% of these two bases were arose among studies. However, it is worth noting that the ratio of G and U does not represent a false-positive rate; instead, it is a false signal caused by various factors. After removing the G and U signals, research has been more concerned with reducing the bias of the A sites to be closer to the background value of the transcriptome [13].

Studies conducted RT-stop and MaP sequencing of specific RNA structures and found that both RT-stop and MaP methods can produce effective single-stranded information, but the information is non-overlapping, speculated that these two methods have the bias of sequence context [34,35]. We calculated the overlapping info of the *E. coli*, *H. sapiens*, and *S. cerevisiae* of transcriptome level and found that two methods had a high of overlapping on average at the transcriptome level (Figure 3C). Transcripts of the same species with similar coverage by two methods were screened. The ratio of overlapping bases of the RT-stop and MaP methods revealed 88.85% and 87.29% in human (143 transcripts) and 98.22% and 95.04% in yeast (2140 transcripts), respectively. From the perspective of data coverage of DMS reactivity, RT-stop had a higher proportion (p < 1e^−4^, Figure 3D, six RT-stop studies, and four MaP studies). This result probably due to the fact that the maximum number of an allowed mismatch for the MaP method is five (and probably the maximum number of the existing alignment software), and there is a limited proportion of mutated bases [16,35], so a higher sequencing depth is needed. The advantage of the MaP method over RT-stop is that it can avoid the modification focusing on the 3’ part of the RNA fragment, regardless of RNA abundance [15].

## DISCUSSION

RSVdb is a comprehensive and user-friendly transcriptome RNA structure database with multiple functions. It not only supports browsing transcriptome RNA structure research studies but also supports online search and prediction of RNA structure with DMS restriction. RSVdb enables users to easily access from multiple platforms through the browser, with additional optimizations for mobile platform access and charting operations.

RSVdb collected and sorted relevant research studies performing DMS probed transcriptome RNA structure, divided them into RT-stop and MaP according to the methods, and then conducted data processing and RNA structure analysis, respectively. On the “Browse” page, the user can obtain statistics of the transcriptome RNA structure studies according to the experimental samples, including sequencing data quality, A/C content, RPKM, DMS reactivity, and Gini coefficient, to provide a general overview of these studies. On the “Viewer” page, users can make full use of data from transcriptome RNA structure studies, including searching for transcripts of interest, obtaining the DMS reactivity of transcripts under different experimental treatments, and predicting RNA structures with DMS restriction. The DMS reactivity data in RSVdb can be downloaded from the “Download” page, and a detailed user manual can be obtained from the “Help” page. Meanwhile, we are committed to improving the user experience, including table and chart support interactive operations and image download, fuzzy search and search suggestion, as well as configuring global CDN services, etc. Also, we open the source code of the front and back ends of the RSVdb in the Github repository, which supports downloading and building the database and achieving the main functions locally.

Although RSVdb has certain advantages in terms of displaying transcriptome RNA structured data, there are still limitations. Currently, the database only contains RNA structure data probed with DMS reagents, but there are still excellent and continuously improving new methods for labeling RNA structures, such as SHAPE and CMCT, etc. In the future, we will continue to update the database, add SHAPE, CMCT, and other methods of detecting the transcriptome RNA structure and further improve the visual display of structural data and website optimization.

## Acknowledgment

The authors would like to thank the Network & Education Technology Center of NWAFU for its support of server hardware and network service. We are grateful to Shuo Gao, Xuanyan Li, Jingjing Son, for their advice on this project in terms of server operation and webpage technology.

## REFERENCES

1. Sharp PA. The Centrality of RNA. Cell 2009; 136:577–580

2. Martin KC, Ephrussi A. mRNA Localization: Gene Expression in the Spatial Dimension. Cell 2009; 136:719–730

3. Mustoe AM, Busan S, Rice GM, et al. Pervasive Regulatory Functions of mRNA Structure Revealed by High-Resolution SHAPE Probing. Cell 2018; 173:181–195 e18

4. Espah Borujeni A, Cetnar D, Farasat I, et al. Precise quantification of translation inhibition by mRNA structures that overlap with the ribosomal footprint in N-terminal coding sequences. Nucleic Acids Res. 2017; 45:5437–5448

5. Faure G, Ogurtsov AY, Shabalina SA, et al. Role of mRNA structure in the control of protein folding. Nucleic Acids Res. 2016; 44:10898–10911

6. Yu H, Meng W, Mao Y, et al. Deciphering the rules of mRNA structure differentiation in Saccharomyces cerevisiae in vivo and in vitro with deep neural networks. RNA Biol 2019; 16:1044–1054

7. Ding Y, Tang Y, Kwok CK, et al. In vivo genome-wide profiling of RNA secondary structure reveals novel regulatory features. Nature 2014; 505:696–700

8. Rouskin S, Zubradt M, Washietl S, et al. Genome-wide probing of RNA structure reveals active unfolding of mRNA structures in vivo. Nature 2014; 505:701–705

9. Hooks KB, Griffiths-Jones S. Conserved RNA structures in the non-canonical Hac1/Xbp1 intron. RNA Biol 2011; 8:552–556

10. Moazed D, Stern S, Noller HF. Rapid chemical probing of conformation in 16 S ribosomal RNA and 30 S ribosomal subunits using primer extension. J Mol Biol 1986; 187:399–416

11. Moazed D, Robertson JM, Noller HF. Interaction of elongation factors EF-G and EF-Tu with a conserved loop in 23S RNA. Nature 1988; 334:362–364

12. Burkhardt DH, Rouskin S, Zhang Y, et al. Operon mRNAs are organized into ORF-centric structures that predict translation efficiency. Elife 2017; 6:

13. Ritchey LE, Su Z, Tang Y, et al. Structure-seq2: sensitive and accurate genome-wide profiling of RNA structure in vivo. Nucleic Acids Res. 2017; 45:e135

14. Incarnato D, Neri F, Anselmi F, et al. Genome-wide profiling of mouse RNA secondary structures reveals key features of the mammalian transcriptome. Genome Biol 2014; 15:491

15. Zubradt M, Gupta P, Persad S, et al. DMS-MaPseq for genome-wide or targeted RNA structure probing in vivo. Nat. Methods 2017; 14:75–82

16. Wang Z, Ma Z, Castillo-Gonzalez C, et al. SWI2/SNF2 ATPase CHR2 remodels pri-miRNAs via Serrate to impede miRNA production. Nature 2018; 557:516–521

17. Guenther UP, Weinberg DE, Zubradt MM, et al. The helicase Ded1p controls use of near-cognate translation initiation codons in 5’ UTRs. Nature 2018; 559:130–134

18. Beaudoin JD, Novoa EM, Vejnar CE, et al. Analyses of mRNA structure dynamics identify embryonic gene regulatory programs. Nat. Struct. Mol. Biol. 2018; 25:677–686

19. Ding YL, Kwok CK, Tang Y, et al. Genome-wide profiling of in vivo RNA structure at single-nucleotide resolution using structure-seq. Nat. Protoc. 2015; 10:1050–1066

20. Deigan KE, Li TW, Mathews DH, et al. Accurate SHAPE-directed RNA structure determination. Proc. Natl. Acad. Sci. U. S. A. 2009; 106:97–102

21. Zubradt M, Gupta P, Persad S, et al. Genome-wide DMS-MaPseq for in vivo RNA structure determination. Nat. Methods 2017; 14:75–82

22. Mortazavi A, Williams BA, McCue K, et al. Mapping and quantifying mammalian transcriptomes by RNA-Seq. Nat. Methods 2008; 5:621–628

23. Cordero P, Kladwang W, Vanlang CC, et al. Quantitative dimethyl sulfate mapping for automated RNA secondary structure inference. Biochemistry 2012; 51:7037–7039

24. Reuter JS, Mathews DH. RNAstructure: software for RNA secondary structure prediction and analysis. BMC Bioinformatics 2010; 11:129

25. Cordero P, Lucks JB, Das R. An RNA Mapping DataBase for curating RNA structure mapping experiments. Bioinformatics 2012; 28:3006–3008

26. Berkowitz ND, Silverman IM, Childress DM, et al. A comprehensive database of high-throughput sequencing-based RNA secondary structure probing data (Structure Surfer). BMC Bioinformatics 2016; 17:215

27. Norris M, Kwok C, Cheema J, et al. FoldAtlas: a repository for genome-wide RNA structure probing data. academic.oup.com

28. Wells SE, Hughes JMX, Haller Igel A, et al. Use of dimethyl sulfate to probe RNA structure in vivo. 2000; 318:479–493

29. Turner DH, Sugimoto N, Freier SM, et al. RNA structure prediction. Annu Rev Biophys Biophys Chem 1988; 17:167–192

30. Mathews DH, Disney MD, Childs JL, et al. Incorporating chemical modification constraints into a dynamic programming algorithm for prediction of RNA secondary structure. Proc. Natl. Acad. Sci. 2004; 101:7287–7292

31. Hauenschild R, Tserovski L, Schmid K, et al. The reverse transcription signature of N-1-methyladenosine in RNA-Seq is sequence dependent. Nucleic Acids Res. 2015; 43:9950–64

32. Roovers M, Wouters J, Bujnicki JM, et al. A primordial RNA modification enzyme: The case of tRNA (m1A) methyltransferase. Nucleic Acids Res. 2004; 32:465–476

33. Lawley PD, Brookes P. Further Studies on the Alkylation of Nucleic Acids and their Constituent Nucleotides. Biochem. J. 1963; 89:127–138

34. Novoa EM, Beaudoin J-D, Giraldez AJ, et al. Best practices for genome-wide RNA structure analysis: combination of mutational profiles and drop-off information. biorxiv 2017;

35. Sexton AN, Wang PY, Rutenberg-Schoenberg M, et al. Interpreting Reverse Transcriptase Termination and Mutation Events for Greater Insight into the Chemical Probing of RNA. Biochemistry 2017; 56:4713–4721

36. Incarnato D, Morandi E, Simon LM, et al. RNA Framework: an all-in-one toolkit for the analysis of RNA structures and post-transcriptional modifications. Nucleic Acids Res. 2018; 46:e97

