## Supplementary material for "RSVdb: A comprehensive database of transcriptome RNA structure": Table 1

Table 1. Studies in RSVdb

| **GEO** | **Class** | **Method** | **Species** | **Samples** | **Transcripts** |
| --- | --- | --- | --- | --- | --- |
| **GSE84537**[15] | MaP | DMS-MaPseq | Drosophila; Human; Yeast | 1;4;10 | 121; 38,052; 51013 |
| **GSE108857**[16] | MaP | DMS–MaPseq | Arabidopsis | 6 | 289 |
| **GSE93959**[17] | MaP | DMS-MaPseq | Yeast | 4 | 13,043 |
| **GSE111962**[36] | MaP | DMS-MaPseq | E.coli | 2 | 5913 |
| **GSE77617**[12] | RT-Stop | DMS-seq | E.coli | 13 | 22,012 |
| **GSE95514**[13] | RT-Stop | Structure-seq2 | Rice | 16 | 11,968 |
| **GSE54106**[14] | RT-Stop | CIRS-seq | Mouse | 2 | 15,676 |
| **SRP027216**[7] | RT-Stop | Structure-seq | Arabidopsis | 4 | 12,702 |
| **GSE45803**[8] | RT-Stop | DMS-seq | Human; Yeast | 11;27 | 232,804; 67,600 |
| **SRP114782**[18] | RT-Stop | DMS-seq | Zebrafish | 78 | 155,032 |
